# *Kentrophoros magnus* sp. nov. (Ciliophora, Karyorelictea), a new flagship species of marine interstitial ciliates

**DOI:** 10.1101/2020.03.19.998534

**Authors:** Brandon K. B. Seah, Jean-Marie Volland, Nikolaus Leisch, Thomas Schwaha, Nicole Dubilier, Harald R. Gruber-Vodicka

**Affiliations:** Max Planck Institute for Marine Microbiology, Celsiusstraße 1, 28359 Bremen, Germany; Department of Limnology and Bio-Oceanography, University of Vienna, Althanstraße 14, A-1090 Vienna, Austria; Department of Integrative Zoology, University of Vienna, Althanstraße 14, A-1090 Vienna, Austria; Max Planck Institute for Developmental Biology, Max-Planck-Ring 5, 72076 Tübingen, Germany; DOE Joint Genome Institute, Lawrence Berkeley National Laboratory, 1 Cyclotron Road, Berkeley, CA 94720, United States of America

**Keywords:** Flagship species, interstitial, meiofauna, pseudotrophosome, symbiosis, taxonomy

## Abstract

The karyorelictean ciliate *Kentrophoros* lacks a defined oral apparatus but has a dense coat of symbiotic bacteria that it consumes by phagocytosis. Body size, shape, and nuclear characters are variable in this genus. We formally describe a new species, *K. magnus* from Elba (Italy), which has unusual folding of its symbiont-bearing surface into pouch-like compartments, a body form that we term “pseudotrophosomal”. *K. magnus* cells are large (2100 ± 700 × 170 ± 23 μm in vivo), but contain only one micronucleus and two macronuclei, although these are much bigger than other *Kentrophoros* (widths 20 ± 2.5 and 31 ± 4.0 μm respectively in *K. magnus*). We also present morphological observations on a close relative from Twin Cayes (Belize), which also has relatively large nuclei (micronuclei 13 ± 1.5 μm, mature macronuclei 20 ± 2.8 μm), but unlike *K. magnus*, it has on average 22 nuclei per cell, with different developmental stages of the macronuclei present simultaneously, and lacks pouch-like folding. Nuclear number and arrangement are important characters for karyorelicts. We suggest the use of a “nuclear formula” to simplify descriptions. Our discovery of large and morphologically distinctive new species underlines the incompleteness of our knowledge about meiofaunal ciliates.

## INTRODUCTION

The karyorelict ciliates are unlike other ciliates in having non-dividing macronuclei with a paradiploid DNA content (Raikov 1985, 1994). Their name derives from an hypothesis that they are a basal lineage of ciliates representing a “primitive” mode of nuclear organization (Corliss and Hartwig 1977). However, molecular phylogenies first using ribosomal RNA (rRNA) genes (Baroin-Tourancheau et al. 1992; Hammerschmidt et al. 1996) and more recently with phylogenomic methods (Gao and Katz 2014) show that they are sister to the heterotrichs, and support the alternative hypothesis that their unusual macronuclei are in fact a derived trait. Almost all known karyorelicts, with the exception of the genus *Loxodes* from freshwater lakes, live interstitially in marine sediments.

*Kentrophoros* Sauerbrey 1928 (syn. *Centrophorus* Kahl 1931, *Centrophorella* Kahl 1935) is a karyorelict genus that is characterized by the lack of an obvious oral apparatus (“mouth”), and a dense coat of ectosymbiotic sulfur bacteria on one side of their cell bodies (Sauerbrey 1928; Kahl 1935; Fenchel and Finlay 1989; Foissner 1995; Seah et al. 2019). The symbiont-bearing surface is non-ciliated, while the other side is covered with somatic kineties; the non-ciliated surface has been homologized with the glabrous stripe of the loxodids and trachelocercids (Foissner 1998); the symbiont-bearing side of *Kentrophoros* would therefore correspond to the “left” side of *Loxodes*. *Kentrophoros* consumes its symbionts by direct phagocytosis through the glabrous stripe, and digestive vacuoles containing bacteria have been observed in the cytoplasm (Raikov 1971; Fenchel and Finlay 1989). When stained with protargol, *K. fistulosus* has condensed anterior kineties associated with fibers that form a “basket-like” structure; these were hypothesized to be oral vestiges (Foissner 1995), however similar structures were not reported in *K. flavus* and *K. gracilis* (Xu et al. 2011). These two studies are the only investigations of *Kentrophoros* to date with silver impregnation methods, although fifteen species have so far been described (Sauerbrey 1928; Fauré-Fremiet 1950, 1951; Dragesco 1954a; b, 1960; Raikov 1962, 1963; Kovaleva 1966; Raikov and Kovaleva 1968; Wright 1982).

Despite the morphological diversity among the different species in this genus, we have previously shown with molecular phylogeny of the 18S rRNA gene that *Kentrophoros* is a monophyletic taxon (Seah et al. 2017). In that study, we discovered a putative morphospecies (designated as morphospecies “H” in that paper) from Elba, Italy which was exceptionally large and had a complex folding of the symbiont-bearing cell surface that forms pouch-like compartments that we named a “pseudotrophosome”. Here, we use live observation, silver impregnation, and electron microscopy to formally describe morphospecies “H” as a new species *K. magnus*. We also present morphological observations on another large morphospecies “FM” from Belize, which is the closest relative to *K. magnus* in the 18S rRNA phylogeny (Seah et al. 2017), and discuss their nuclear characters and infraciliature.

## MATERIALS AND METHODS

### Collection of specimens

*Kentrophoros magnus* was collected in July and November 2014 from the island of Elba, Italy, at the bays of Cavoli (42.734192 °N, 10.185868 °E, 12.8 m depth) and Sant’ Andrea (42.808561 °N, 10.142275 °E, 7.3 m depth). Sandy sediment was gently shoveled by Scuba divers into sample containers. Ciliates were extracted by gentle stirring of sediment and decantation with seawater into trays for sorting. *Kentrophoros* sp. “FM*”* was collected in June/July 2015 from Twin Cayes, Belize. Sediment off the southern end of the island (16.82356°N, 88.106150°W, 1.5 m depth) was scooped into a plastic bucket. Ciliates were extracted by both decantation and the seawater ice method (Uhlig 1964).

### Pyridinated silver impregnation

Silver impregnation followed the Fernández-Galiano method (Fernández-Galiano 1976) with some modifications (Ma et al. 2003; Foissner 2014). Solutions were prepared according to Fernandez-Galiano (1976), except that Bacto Tryptone (BD Biosciences) was substituted for peptone. Fernández-Galiano fluid was prepared freshly each day from pyridine (15 μl), Rio-Hortega solution (175 μl), tryptone (40 μl), and deionized water (800 μl). Live *Kentrophoros* individuals were transferred into ca. 50 μl drops of seawater on glass slides (Superfrost Plus, Thermo Scientific), fixed with 50 μl of 4% (w/v) formaldehyde (Electron Microscopy Sciences) in seawater for 1-3 min, rinsed three times in deionized water, and then fixed for a further 1-3 min with 50 μl of 4% formaldehyde in deionized water. 50 μl of Fernández-Galiano fluid was added to the drop, mixed gently, and incubated for 1 min at room temperature. The slide was then warmed on a hotplate at 58 ºC with shaking (250 rpm), and removed when the drop turned golden brown. After removing excess fluid, the specimen was gently rinsed with water and mounted with a coverslip for photomicrography with bright-field illumination with either an Olympus BX53 or Zeiss Axioskop microscope. For permanent preparations, the coverslip was removed, the preparation was fixed with 50 μl of 2.5% (w/v) sodium thiosulfate (Sigma-Aldrich) for 5 min at room temperature, and rinsed three times with water, before being air-dried, dehydrated through a series of ethanol (70, 80, 96% v/v), infiltrated with a 1:1 mixture of ethanol and Roti-Histol (Carl Roth) followed by 100% Roti-Histol, and mounted with Permount (Fisher Scientific) under a coverslip.

It was impractical to count the number of ciliary rows directly in *K. magnus*, because of the large size, folding, and distortion of impregnated specimens. Instead, the kinety spacing for each specimen was estimated by 3 measurements of the spacing of between 5 and 10 kineties in the mid-body region. The number of ciliary rows at mid-body was estimated as the ratio of twice the mid-body width to the mean inter-ciliary-row spacing.

### Scanning electron microscopy (SEM)

Samples for SEM were fixed at 4 ºC with 2.5% (w/v) glutaraldehyde buffered with 3× PHEM (120 mM PIPES, 50 mM HEPES, 20 mM EGTA, 4 mM magnesium chloride, all from Sigma-Aldrich) (Montanaro et al. 2016) for >10 h, then with a mixture of 2% glutaraldehyde (Electron Microscopy Sciences) and 2% formaldehyde buffered with 2× PHEM for >10 h, and then washed three times in 4.5× PHEM, and stored in washing buffer at 4 ºC until use. Fixed samples were post-fixed in 1% (w/v) osmium tetroxide aq. (Electron Microscopy Sciences) for 2 h, and then washed at least 3 times with water. Cells were settled on glass coverslips coated with poly-L-lysine, dehydrated in ethanol (30, 40, 50, 60, 70, 80, 90, 96, 96, 100, 100%, > 20 min per step), critical-point-dried with carbon dioxide on an EM CPD300 machine (Leica), and imaged on a Quanta FEG 250 scanning electron microscope (FEI) with 1 to 3 kV electron beam and secondary electron detector.

### Transmission electron microscopy (TEM)

Samples for TEM were fixed in 1% (w/v) osmium tetroxide with 0.1 M sodium cacodylate, 1,100 mOsm liter^−1^, pH 7.4 (Electron Microscopy Sciences) for 2 h, washed three times in 0.1 M sodium cacodylate, 1,100 mOsm liter^−1^, pH 7.4 overnight (> 12 h), washed three times with distilled water, dehydrated as for SEM (above), and embedded in EMBed 812 resin (Electron Microscopy Sciences) using acetone as an intermediate solvent. The EMBed 812 was mixed in the “hard” formulation and cured at 60 ºC for 24 h. Ultrathin sections transferred to slot grids with a Formvar support film were stained with 0.5% (w/v) uranyl acetate and 3% (w/v) lead citrate before examination with a Zeiss Libra 120 transmission electron microscope operating at 120 kV.

## RESULTS

### Description of *Kentrophoros magnus*

NB: Full morphometric data are given in Table 1, and are not repeated in the description.

**Table 1.** Morphometric characters of *K. magnus* from Cavoli (CAV) and Sant’ Andrea (SAN) populations

| Character | Population | Treatment | Mean | SD | Min | Max | CV (%) | *n* |
| --- | --- | --- | --- | --- | --- | --- | --- | --- |
| Body length (µm) | SAN | Live | 1930 | 604 | 1030 | 3370 | 31.3 | 19 |
|  | CAV | Live | 2110 | 696 | 931 | 3450 | 32.9 | 16 |
|  | CAV | Silver | 1418 | 481 | 647 | 2791 | 33.9 | 33 |
| Body width (µm) | SAN | Live | 164 | 31.9 | 113 | 217 | 19.4 | 19 |
|  | CAV | Live | 171 | 23.3 | 142 | 212 | 13.6 | 16 |
|  | CAV | Silver | 212 | 47.3 | 151 | 396 | 22.4 | 33 |
| Head length (µm) | SAN | Live | 263 | 103 | 100 | 470 | 39.1 | 19 |
|  | CAV | Live | 320 | 127 | 119 | 545 | 39.7 | 16 |
|  | CAV | Silver | 208 | 45.0 | 105 | 274 | 21.6 | 31 |
| Head width (µm) | SAN | Live | 75.3 | 14.8 | 47.9 | 104 | 19.6 | 19 |
|  | CAV | Live | 84.6 | 13.4 | 66.6 | 116 | 15.9 | 15 |
|  | CAV | Silver | 77.2 | 12.9 | 56.2 | 111 | 16.7 | 29 |
| Tail length (µm) | SAN | Live | 116 | 37.7 | 61.4 | 175 | 32.6 | 15 |
|  | CAV | Live | 95.8 | 33.4 | 44.2 | 138 | 34.8 | 8 |
|  | CAV | Silver | 93.4 | 31.1 | 43.3 | 167 | 33.3 | 26 |
| Ciliary row spacing (µm) | CAV | Silver | 3.0 | 0.55 | 2.0 | 4.1 | 18.3 | 22 (66) |
| No. ciliary rows at mid body (est.) | CAV | Silver | 140 | 25.3 | 108 | 210 | 18.0 | 20 |
| MIC width (µm) | CAV | Silver | 19.6 | 2.54 | 16.4 | 24.6 | 13.0 | 12 (12) |
| MAC width (µm) | CAV | Silver | 30.5 | 4.03 | 25.2 | 39.5 | 13.2 | 10 (19) |

<sup>a</sup> *n* indicates number of individual cells measured. Where more than one measurement was made per cell, total number of measurements are given in brackets.

#### Shape and behavior

Large and vermiform ciliates, with club-shaped anterior (“head”) and pointed posterior (“tail”) regions (terminology of anatomical orientation in Fig. 1) Cell body appears bright white under oblique dark-ground illumination because of light scattered by the ectosymbiotic sulfur bacteria. Symbiont-bearing regions appear dark brown or black under transmitted illumination. Ectosymbiont coat absent from hyaline head and tail regions. Head slightly hooded and asymmetrical, with the right side more curved than the left side. (Fig. 2, 3)

**Fig. 1.**
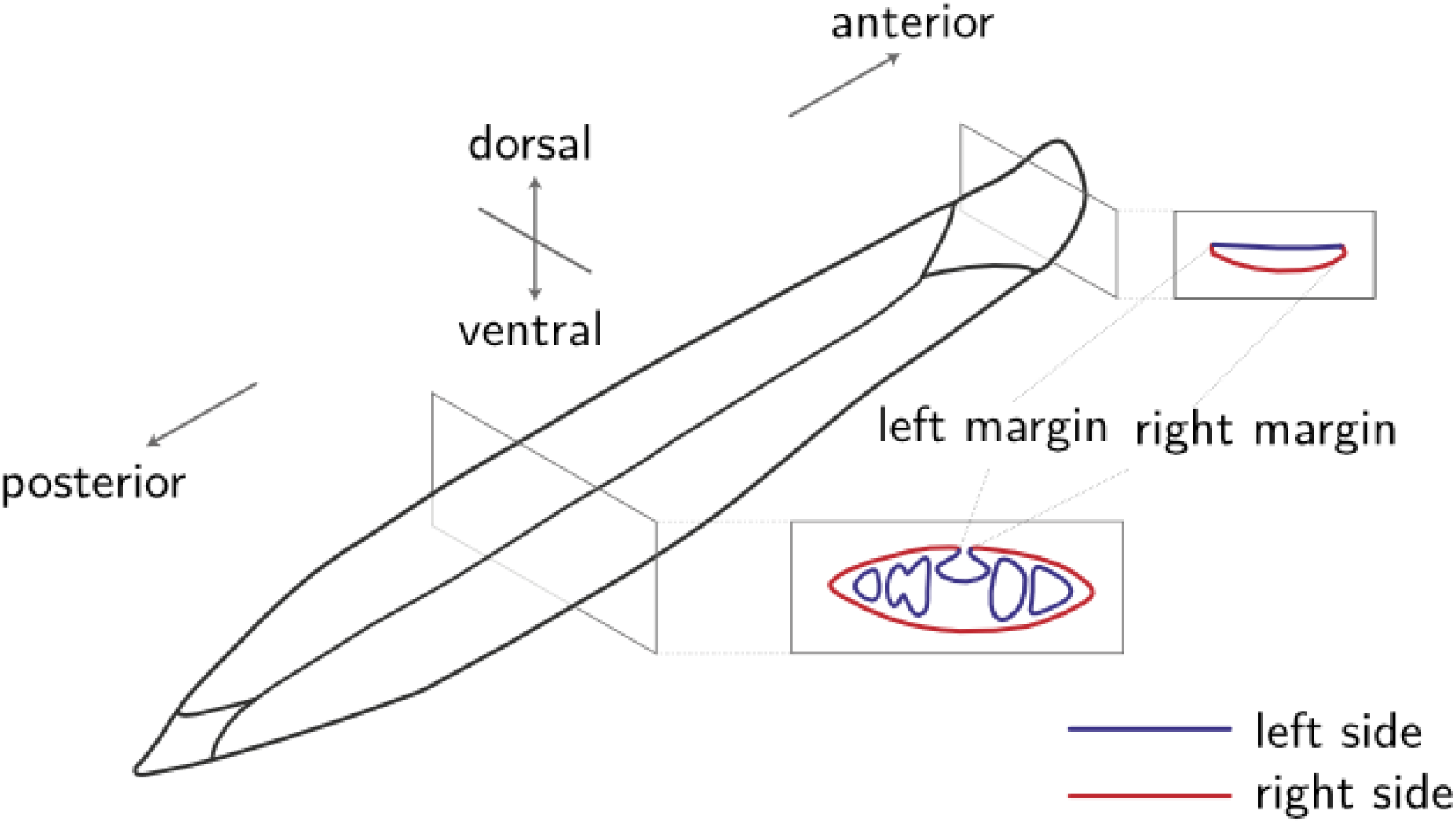
Terminology of anatomical orientation in *K. magnus*

**Fig. 2.**
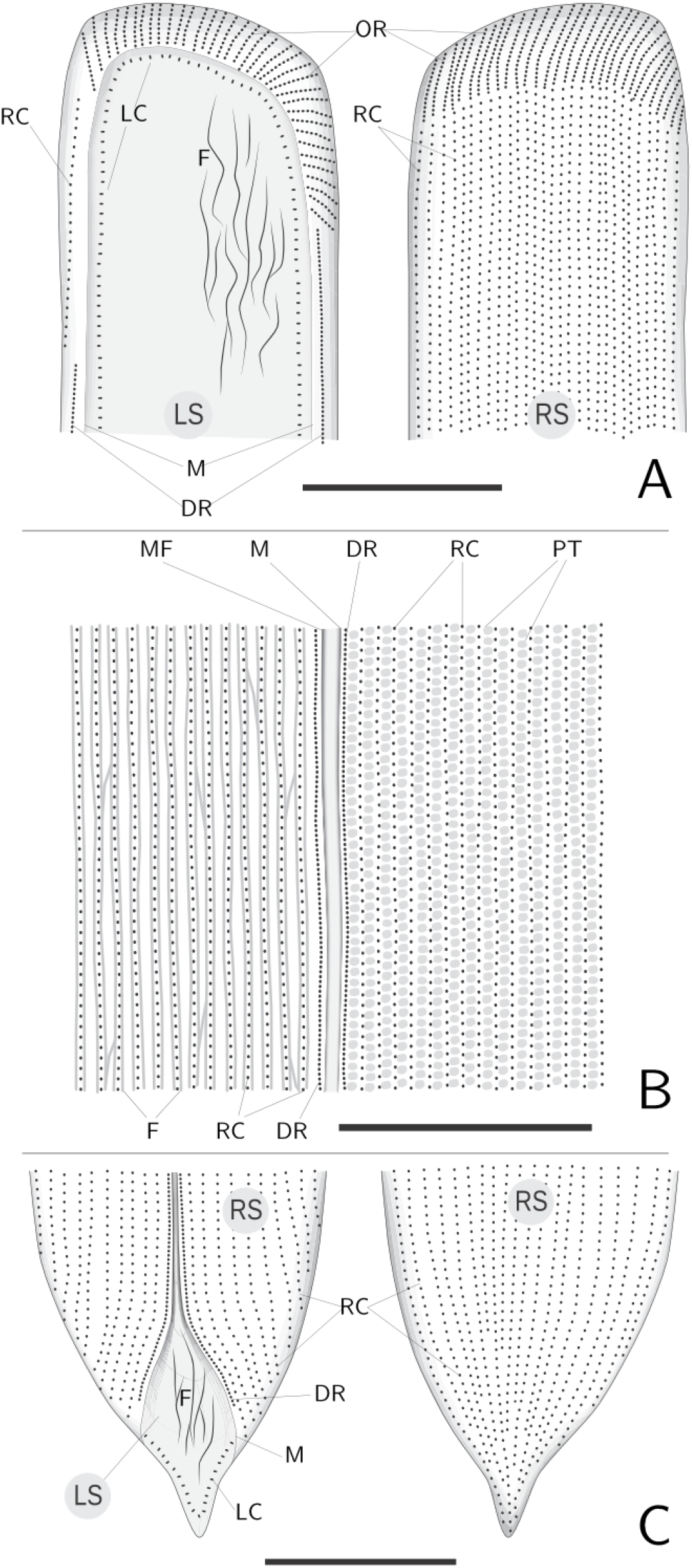
Diagram of infraciliature of *K. magnus*. **A**. Head. **B**. Mid-body. **C**. Tail. Legend: DR, Dense right ciliary row; F, Fiber; LC, Left ciliary row; LS, Left side; M, Cell anatomical margin; MF, Median furrow; OR, Oblique right ciliary row; PT, Protrichocyst; RC, Right somatic ciliary row; RS, Right side. Scales: 25 μm.

**Fig. 3.**
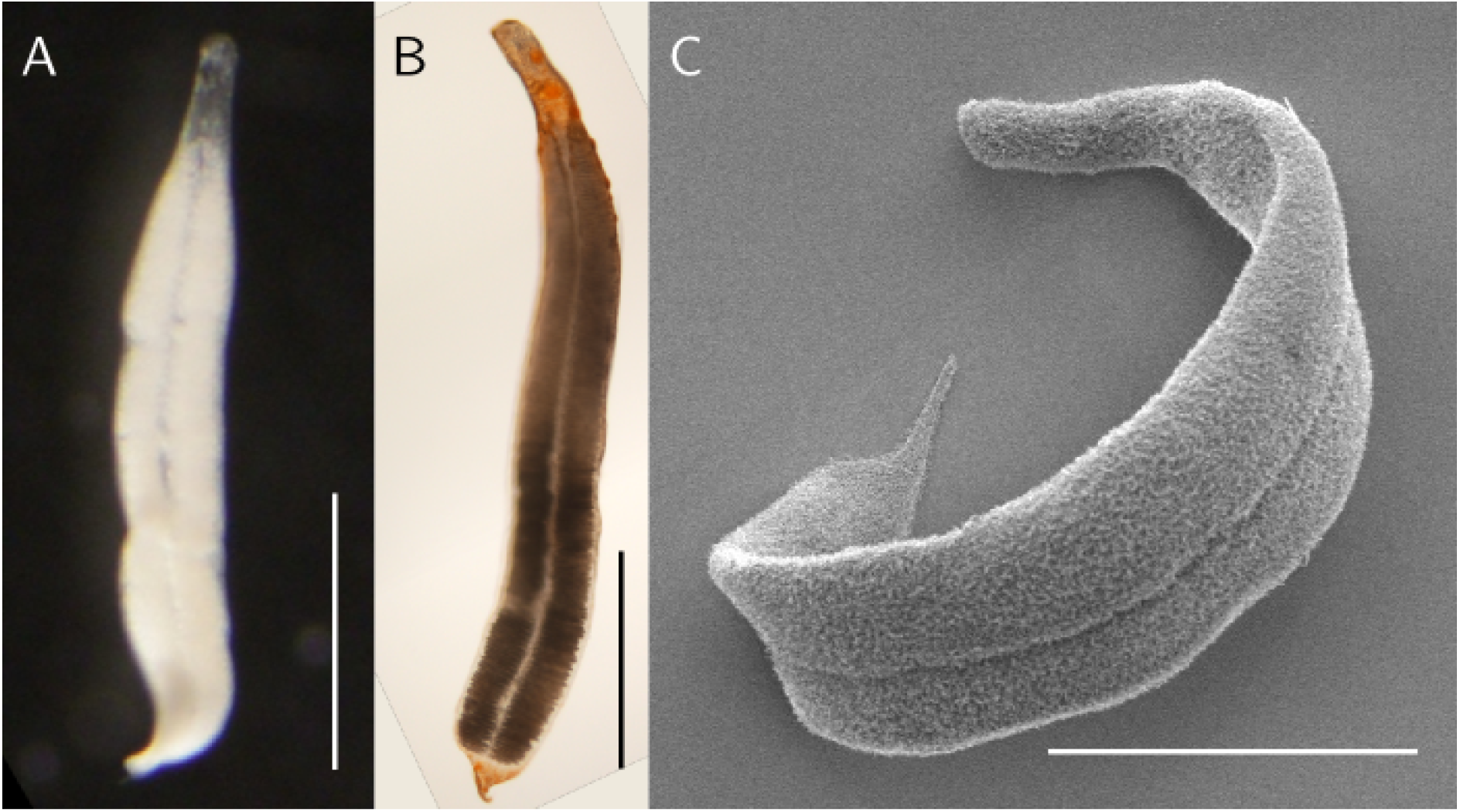
Habitus of *K. magnus*. **A**. Live, dorsal view, incident light. White color is due to light scattered by ectosymbionts. Scale: 500 μm. **B**. Silver-impregnated, dorsal view, brightfield. Scale: 500 μm. **C**. SEM. Scale: 250 μm.

Entire body involuted except for head and tail, with ciliated, anatomically ventral side (sensu (Fauré-Fremiet 1950b)) on outer surface of the folded body, and symbiont-bearing dorsal side on inside surface. Previous descriptions of *Kentrophoros* have homologized the dorsal (symbiont-bearing) and ventral surfaces with the left and right sides respectively of the loxodid ciliates (Foissner 1995, 1998; Xu et al. 2011). For consistency, we reserve the terms “left” and “right” to refer to the anatomical surfaces, whereas “dorsal” and “ventral” are for gross orientation only. For example, in the involuted body region, the right side of the ciliate is exposed on both the dorsal and ventral surfaces. (Fig. 1)

Anatomical margins meet at a median suture (Fig. 1, 6). Ciliate cytoplasm is thicker along anterior-posterior axis below median suture, forming a medial cytoplasmic strand (Fig. 6). Unlike in *K. fistulosus* (Foissner 1995), we have not observed any individuals of this species that survive the unfolding of the symbiont-bearing surface to the outside.

Movement is by slow gliding with the head forward, though sometimes also backwards. In Petri dishes, the ciliates glide along the bottom in straight lines or in circles, sometimes corkscrewing or performing rolls. Upon disturbance (e.g. shaking of dish) they frequently contract and coil up, or briefly freeze and change direction. Movement after disturbance usually begins with the head, with the rest of the body straightening out behind it.

Damaged individuals, which have lost their heads or tails or have suffered some other injury, can heal and regenerate missing parts within hours to days when stored in sediment.

#### Pseudotrophosome

Bacterial symbionts appear packed into regular series of pouches projecting laterally from the median axis (Fig. 4F), with monolayer on median thickening bridging the two sides (Fig. 6). Pouches formed by folds and undulations of left side, not topologically inside the host body, but still contiguous with external medium. Lateral edges of folded ciliate body, distal to pseudotrophosome, form hyaline “wings” (Fig. 6).

**Fig. 4.**
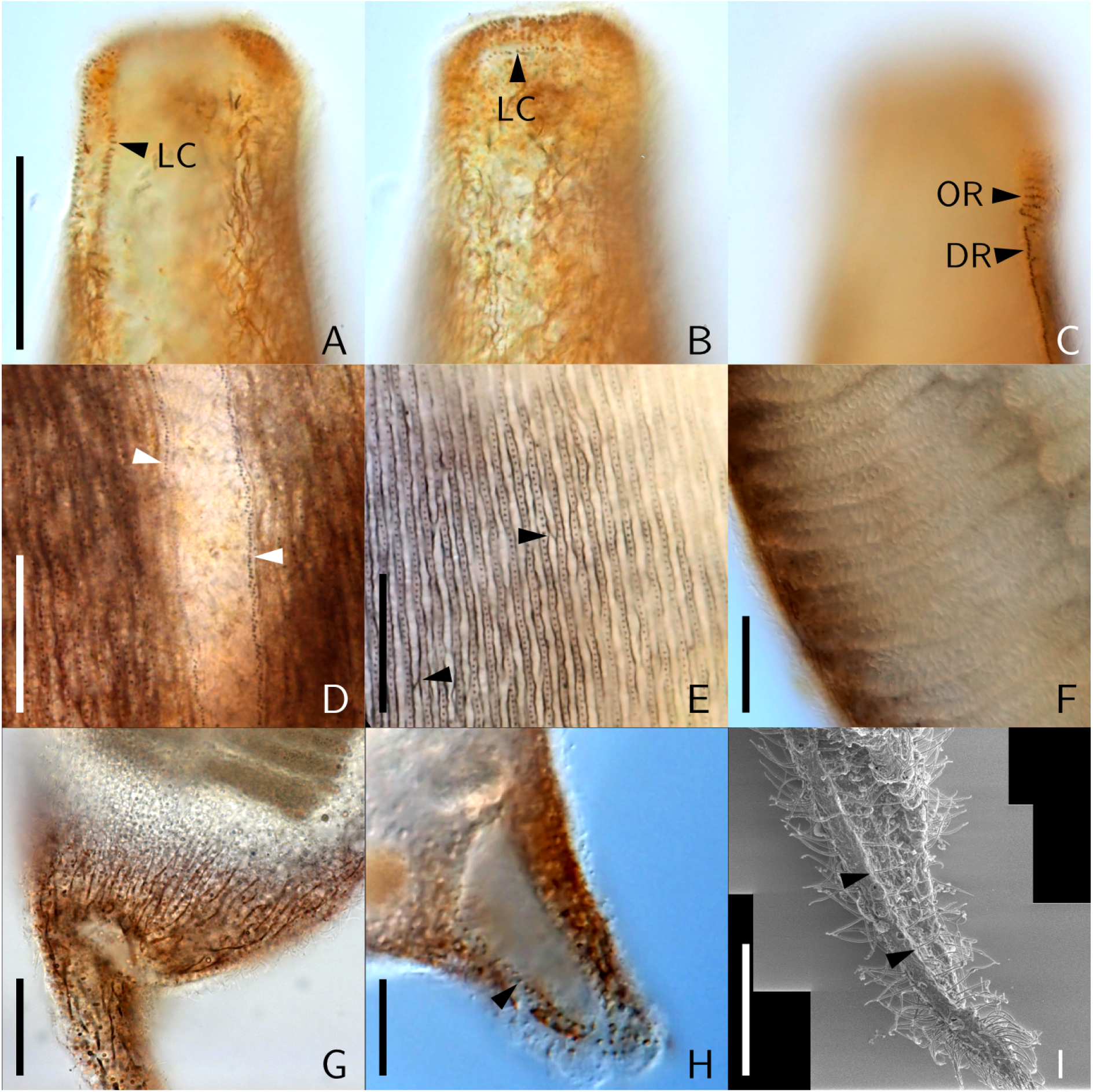
Infraciliature of *K. magnus* in silver-impregnated specimens under brightfield illumination (A to H) and SEM (I). **A**to **C**. Anterior region from same specimen in different focal planes. DR, dense right ciliary row; LC, left ciliary row; OR, oblique right ciliary rows. Scale 50 μm. **D**. Midbody region, dense ciliary rows (arrowheads) flanking median furrow. Scale: 25 μm. **E**. Right somatic ciliary rows and fibers at midbody, with fibers crossing between rows (arrowheads). Scale: 25 μm. **F**. Symbiont “pouches” at midbody. Scale: 25 μm. **G**. Posterior region, branching fibers. Scale: 25 μm. **H**. Left ciliary row (arrowhead) at posterior region. Scale: 15 μm. **I**. Left ciliary row (arrowheads), SEM. Scale: 15 μm.

**Fig. 5.**
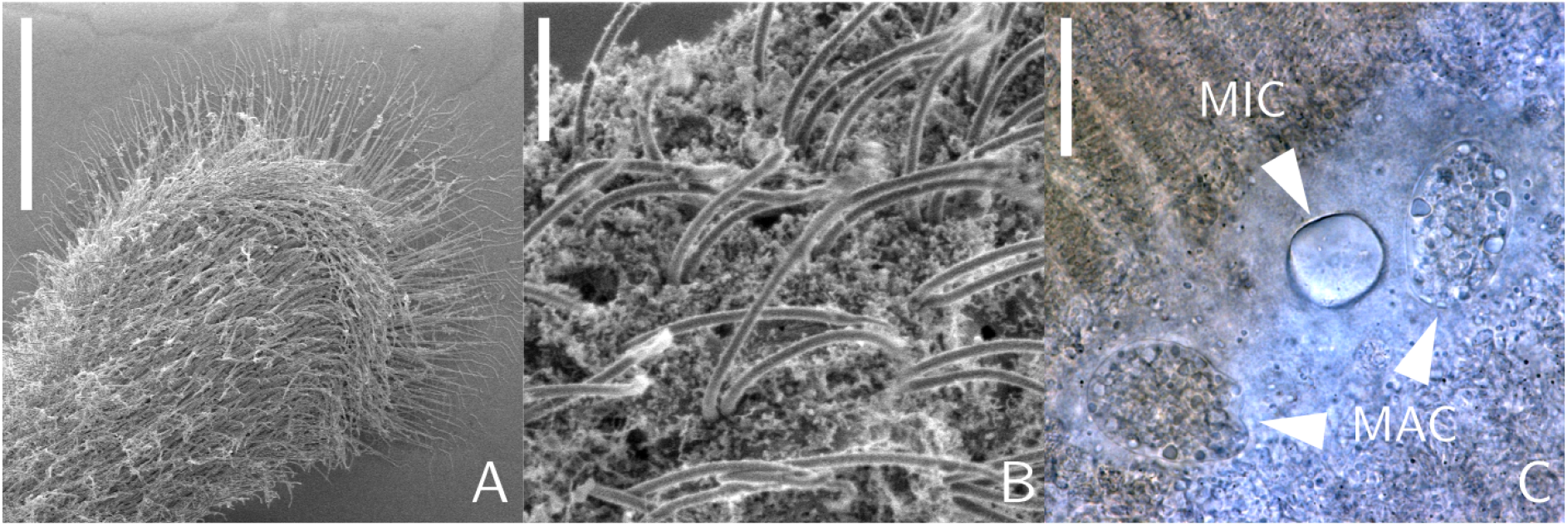
Detailed views of *K. magnus*. **A**. Ventral view of anterior portion, SEM. Scale: 25 μm. **B**. Paired somatic cilia on right side close to head region, SEM. Scale: 2 μm. **C**. Nuclear group comprising micronucleus (MIC) and two macronuclei (MAC) at mid-body, brightfield. Scale: 25 μm.

**Fig. 6.**
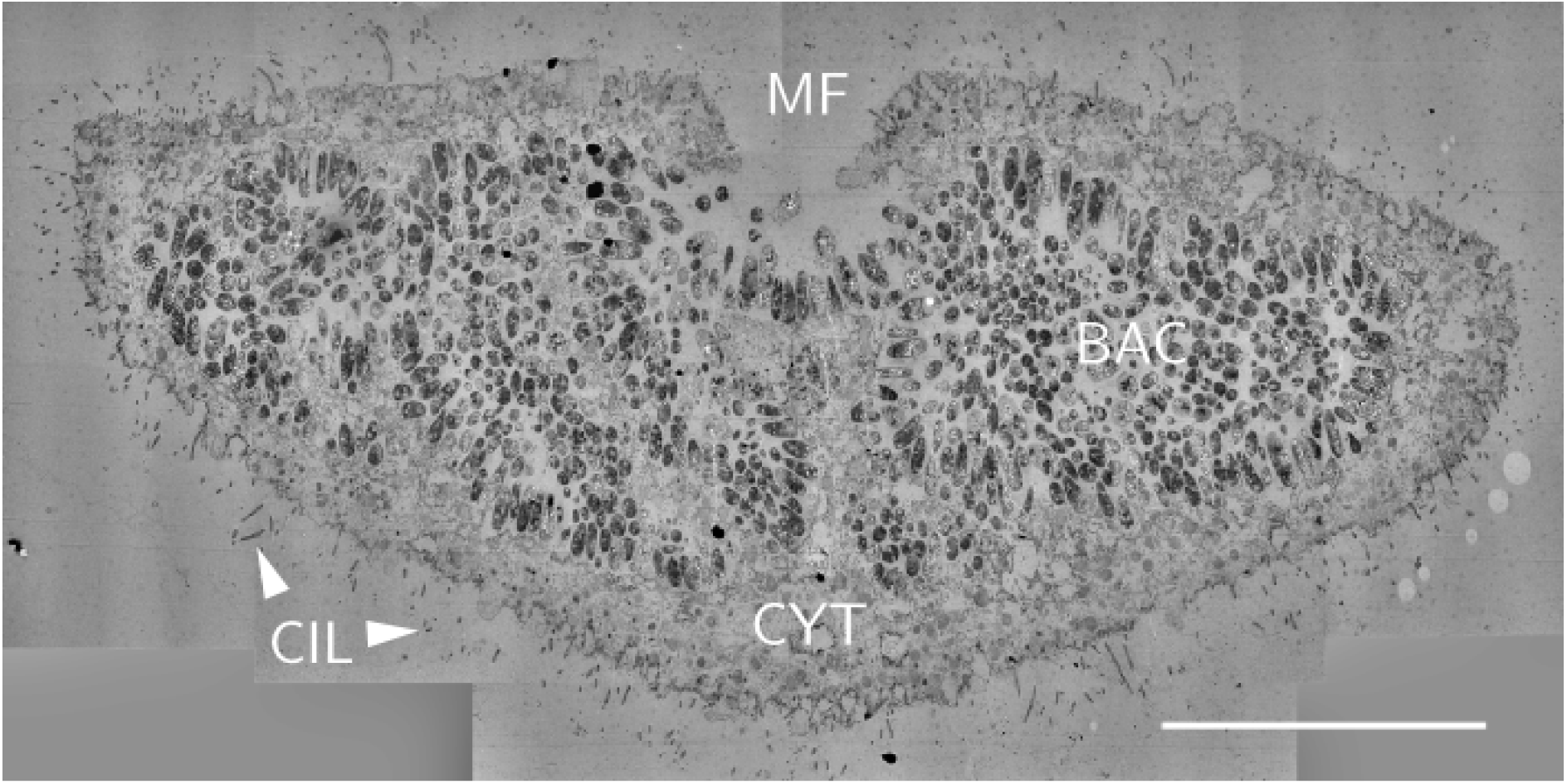
Cross section of *K. magnus* at mid-body, TEM composite image. BAC, bacterial symbionts; CIL, cilia; CYT, ciliate cytoplasm; MF, Median furrow. Scale: 25 μm.

#### Infraciliature

Right side with somatic kineties, parallel to main body axis. Right kineties spaced 3.0 ± 0.55 μm apart in the mid-body region, i.e. about 100 to 200 kineties at mid-body. Ciliary rows become obliquely oriented and with kinetids more densely arranged towards anterior end of head. (Fig. 2, 5A) Close to each margin is a single kinety with densely-arranged kinetids, flanking the median suture of involuted body region, extending into head and terminating before the obliquely-oriented anterior ciliary rows, but not extending into tail (Fig. 4D).

Individual kinetosomes could not be distinguished in silvered specimens; somatic kinetids in head and tail were inferred to be diciliated dikinetids because cilia were seen to emerge in pairs in SEM micrographs (Fig. 5B) and in differential interference contrast, but midbody somatic kineties were too closely packed to unambiguously state whether they are mono- or diciliated.

Left side with monociliated kinety near margin that apparently curves around in a loop at head and at tail (Fig. 2, 4A, 4B). Kinetids of left kinety associated with short rod-like silver-impregnated extensions that may be fibers. Right kinetids lack such projections and appear dot-like when silvered. Left kinety observed in head and tail regions, but could only be followed for a short distance into the mid-body region because of the involution and thickness of the cell.

Each right kinety in mid-body region flanked by pair of silver-impregnated fibers, neither of which is more strongly stained than the other (Fig. 4E). Fibers run largely longitudinally, but cross over between rows at irregular intervals, forming an interconnected network. Additional irregularly-branched fibers present in head and tail region, that also ramify into the cytoplasm (Fig. 4A, 4G).

Effectiveness of silver impregnation was consistently stronger in the head and tail regions, which are symbiont-free, than in the mid-body region over the pseudotrophosome (Fig. 4F). In a few specimens where symbionts were absent from parts of the involuted body region, staining intensity was stronger in symbiont-free parts.

#### Nuclei

Nuclei grouped in a single row within medial cytoplasmic strand, roughly at midpoint of cell. The group comprises one micronucleus (MIC) flanked by one or two macronuclei (MAC) (Fig. 5C). Nuclei strongly refractile under brightfield illumination, but not stained by Fernández-Galiano method. MIC oblate spheroid, width 19.6 ± 2.5 μm, appear internally uniform in both brightfield illumination and in DAPI-stained semithin sections. MAC ovoid to teardrop-shaped in outline, width 30.5 ± 4.0 μm, appear internally heterogeneous in brightfield. In stained semithin sections, MAC contain strongly-stained chromocenters interspersed among a more weakly-stained background.

#### Cytoplasmic inclusions

Clear refractile bodies form regular rows between right kineties. In cross-sections, they stain with toluidine blue and are brightly autofluorescent under excitation wavelengths ranging from ultraviolet to green light.

#### Comparison to other species

This is the only known *Kentrophoros* species with a well-developed pseudotrophosome; other species may be involuted, but lack the pouch-like foldings of the symbiont layer. In terms of nuclear size, it is most similar to *K.* sp. “FM” and “G” (see below), but in the number and arrangement of nuclei, it resembles *K. fasciolatus*, which also has two macronuclei intercalated by one micronucleus.

#### *Comparison of Cavoli and Sant’ Andrea populations of* K. magnus

*K. magnus* collected in two localities in Elba – Cavoli and Sant’ Andrea – had the same qualitative characters, and overlapped in their morphometric characters. A multivariate analysis of variance (MANOVA) test on live specimen morphometrics (excluding tail length, because of missing data) for the two populations did not reject the null hypothesis of equal means between localities (*p* = 0.376, *n* = 34, Pillai-Bartlett statistic). Individual Welch two-sample *t*-tests on the parameters body width (*p* = 0.512, *n* = 35) and head width (*p* = 0.0644, *n* = 34) also did not reject the null hypothesis. 18S rRNA sequences of both were also virtually identical, with 100% identity to each other, except one Cavoli sequence, which had 3 nucleotide substitutions (out of 1557 bp) vs. the other sequences. We therefore regard these two populations as conspecific.

### Observations on *Kentrophoros* sp. “FM”

Two morphospecies, “FM” and “G”, from Twin Cayes (Belize) were found to be the closest relatives to *K. magnus* in the 18S rRNA phylogeny (Seah et al. 2017). The material was inadequate for a formal description, because the infraciliature was poorly stained, but we present morphological observations on morphospecies “FM” for comparison with *K. magnus*. (NB: Full morphometric data are given in Table 2, and are not repeated in the description.)

**Table 2.** Morphometric characters of *K.* sp. “FM”

| **Character** | **Treatment** | **Mean** | **SD** | **Min** | **Max** | **CV (%)** | **n<sup>a</sup>** |
| --- | --- | --- | --- | --- | --- | --- | --- |
| Body length (µm) | Live | 1130 | 329 | 596 | 1720 | 29.1 | 14 |
|  | Silver | 1030 | 261 | 716 | 1490 | 25.3 | 14 |
| Body width (µm) | Live | 77.2 | 14.1 | 52.7 | 99.8 | 18.3 | 14 |
| Ciliary row spacing (µm) | Silver | 3.79 | 1.11 | 2.64 | 6.37 | 29.3 | 9 |
| MIC width (µm) | Silver | 12.8 | 1.47 | 9.3 | 16.6 | 11.5 | 15 (69) |
| MAC (1)<sup>b</sup> width (µm) | Silver | 20.2 | 2.78 | 13.8 | 26.6 | 13.8 | 8 (28) |
| MAC (2)<sup>b</sup> width (µm) | Silver | 13.7 | 3.33 | 8.6 | 23.2 | 24.3 | 13 (43) |
| MAC (3)<sup>b</sup> width (µm) | Silver | 8.10 | 1.10 | 6.8 | 11.4 | 13.6 | 12 (27) |
| Number of MIC | Silver | 5.93 | 1.71 | 4 | 10 | 28.8 | 15 |
| MAC:MIC ratio | Silver | 2.71 | 0.66 | 2 | 4 | 24.5 | 15 |

<sup>a</sup> Number of individual cells from which measurements were made; where different, number of actual measurements given in brackets.
<sup>b</sup> Numbers indicate different putative developmental stages of macronuclei, described in text.

#### Shape and behavior

Large and ribbon-like ciliates; head and tail tapered, hyaline (Fig. 7). Left side covered by dense coat of rod-shaped symbiotic bacteria, except for head and tail ends. Margins undulating in vivo, cell body otherwise not involuted or folded. Cytoplasm thicker along anterior-posterior midline, forming medial cytoplasmic strand. Cells with silvery appearance under incident light in vivo, with apparently clearer stripe down midline corresponding to medial cytoplasmic strand.

**Fig. 7.**
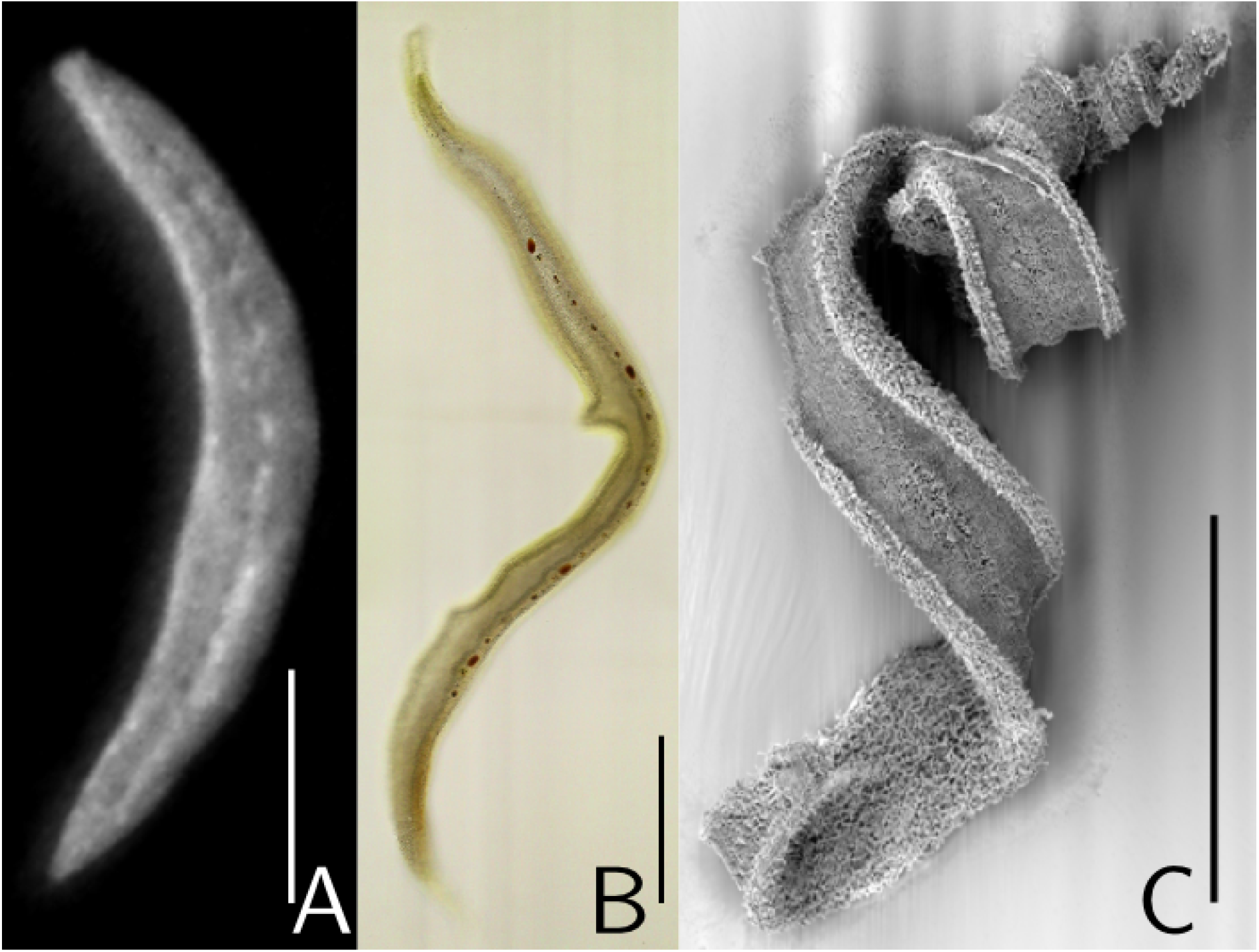
Habitus of *K.* sp. “FM”. **A**. Live, incident light. **B**. Silver-impregnated, brightfield. **C**. SEM. (Scales 150 μm)

#### Infraciliature

Right side with longitudinal somatic kinety rows, but silver impregnation poor. Somatic kineties inferred to be diciliated dikinetids because cilia seen to emerge in pairs under SEM (Fig. 9A). Densely-arranged kineties not observed. Left side with monociliated kinety directly adjacent to bacteria at margins (Fig. 9B).

**Fig. 8.**
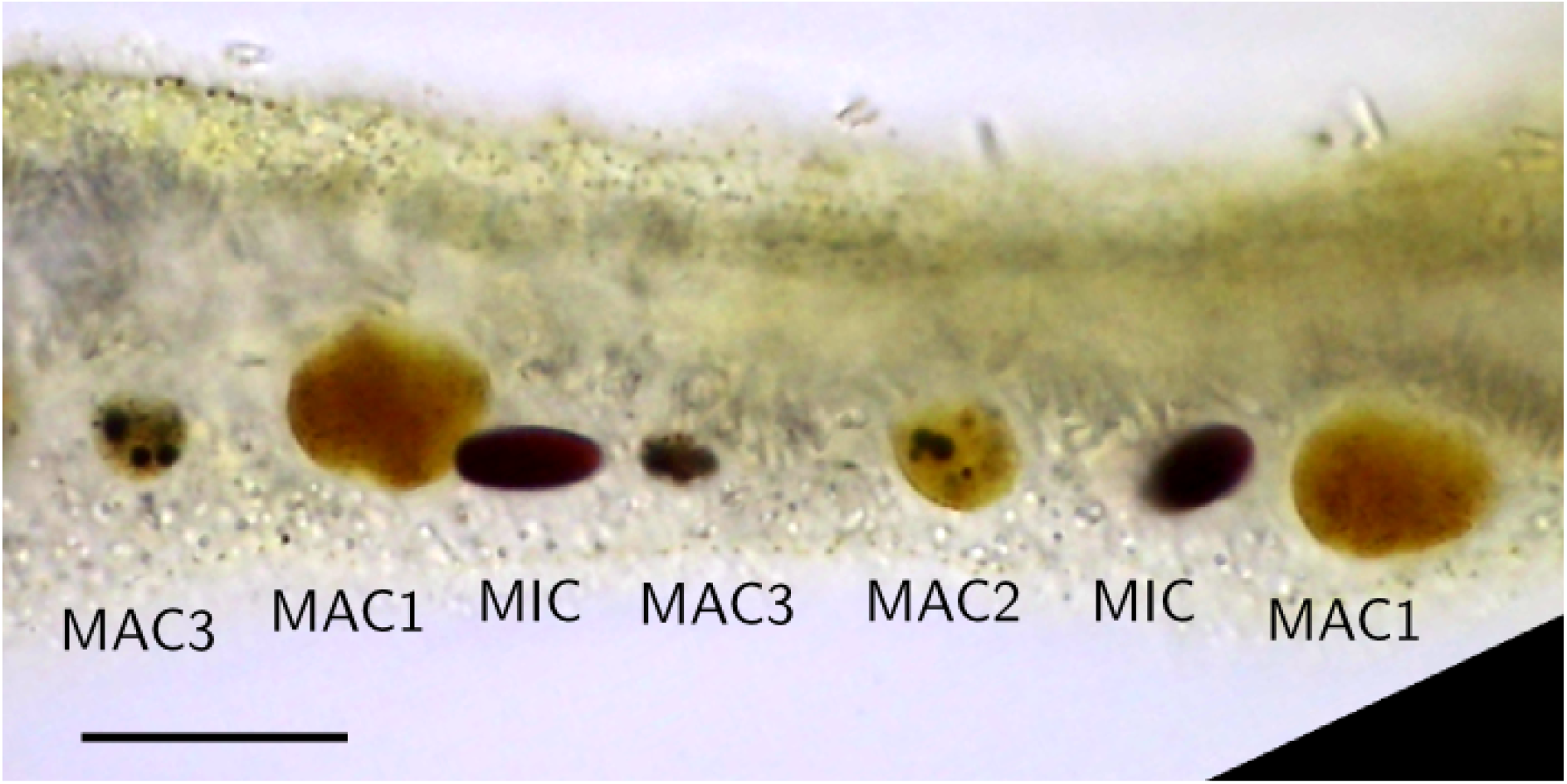
Detailed view of *K.* sp. “FM” nuclei, showing MIC and different MAC stages. Silver-impregnated, brightfield. Scale: 50 μm.

**Fig. 9.**
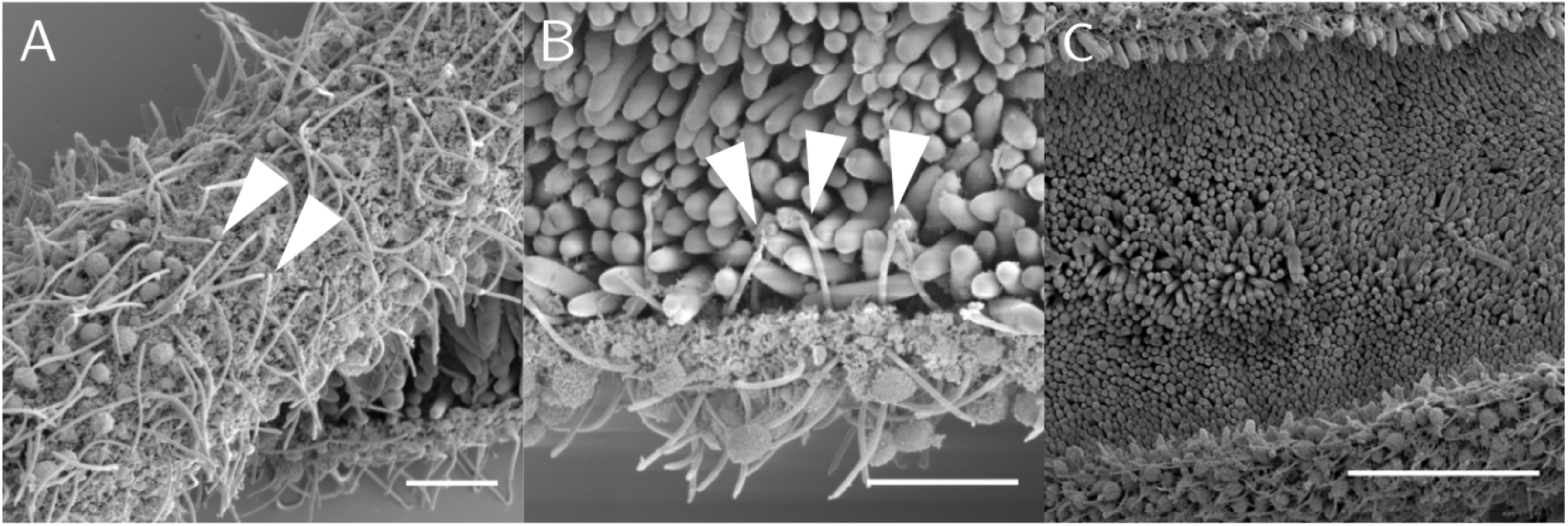
Detail of *K.* sp. “FM” under SEM. **A**. Paired somatic cilia on right side (arrowheads). Right side covered in fluffy material that may be mucus. Scale: 5 μm. **B**. Monociliated left ciliary row (arrowheads) directly adjacent to rod-shaped bacteria. Scale: 5 μm. **C**. Bacterial coat on left side at midbody. Scale: 20 μm.

#### Nuclei

Nuclei arranged in single irregularly-spaced row in median cytoplasmic strand, well-stained by pyridinated silver method. MIC interspersed by varying numbers of MAC (Fig. 8). MIC oblate spheroidal, dense and homogeneously stained. MAC of three types: (1) spheroidal, larger than MIC, without dense spheroidal chromocenters, (2) spheroidal, similar-sized to MIC, with dense spherical chromocenters, (3) irregular outline, smaller than MIC, most of volume comprising dense chromocenters. Type (2) most commonly observed, but all three MAC types can be present in same cell. MIC usually directly flanked by type (1) or (2) MAC.

#### Comparison to other species

*K.* sp. “FM” is similar in size and nuclear characters, and has nearly identical 18S rRNA sequence to another morphospecies from same locality, designated as “G” (Seah et al. 2017). However, unlike the open body form of *K.* sp. “FM”, *K*. sp. “G” has an involuted body shape with pouch-like folding of the symbiont-bearing surface, although not as convoluted or pronounced as the pseudotrophosome in *K. magnus*.

## DISCUSSION

### Nuclei

The nuclei of *K. magnus* and *K.* sp. “FM” are larger than any *Kentrophoros* or karyorelict species thus far described (Raikov 1985; Carey 1992), which have typical sizes of 1-2 μm (MIC) and 4-6 μm (MAC) (Table 3). In the 18S rRNA phylogeny (Seah et al. 2017), *K. magnus* (designated as “H”) and *K.* sp. “FM” belong to the same clade within *Kentrophoros* but the branches within this clade are not well resolved. It is therefore not clear if the enlarged nuclei are a shared derived trait. We speculate that these large nuclei could have originated from a polyploidy event, which could be tested if sequences from nuclear loci are available.

**Table 3.** Sizes of macro-(MAC) and micronuclei (MIC) of *Kentrophoros,* reported or depicted in illustrations in published literature

| **Species** | **Treatment** | **MIC width (µm)** | **MAC width (µm)** | **Reference** |
| --- | --- | --- | --- | --- |
| *K. magnus* | Silver carbonate | 19.6 | 30.5 | This study |
| *K. sp. "FM"* | Silver carbonate | 12.8 | 13.7 (mature stage) | This study |
| *K. canalis* | Fixed | 2 | 4-6 | Wright 1982 |
| *K. fistulosus* | Protargol | 2.1 | 4.1 | Foissner 1995 |
| *K. flavus* | Feulgen | 1-1.2 | 2 | Raikov and Kovaleva 1996 |
| *K. flavus* | Protargol | not reported | 2-4 | Xu et al. 2011 |
| *K. graciles* | Feulgen | not reported | 3-4 | Raikov 1963 |
| *K. lanceolatus* | not reported | not reported | 4 | Fauré-Fremiet 1951 |

*K. magnus* has only a single cluster of nuclei (2 MAC + 1 MIC) despite being such a large ciliate cell. This is puzzling, because other millimeter-scale *Kentrophoros*, such as *K. fistulosus*, have multiple groups or clusters of nuclei in a longitudinal row. Most cell fragments produced by mechanical injury, in the case of *K. fistulosus*, would therefore be nucleated and capable of regeneration to whole individuals. This, presumably, is also a means of asexual reproduction. For *K. magnus*, however, because the nuclei are few and clustered in a small central region of the cell, fragments are more likely to be anucleate and therefore incapable of reproduction.

The different macronucleus morphologies of *K.* sp. “FM” appear to correspond to developmental stages that have been described for other karyorelicts (Raikov 1985). We interpret the three observed types as: (1) swollen macronuclear anlage with vacuolar/granular contents but without condensed chromocenters, (2) mature macronucleus with prominent chromocenters and nucleoli, (3) degenerate, senescent nuclei. This interpretation is supported by the appearance of stained macronuclei, and the fact that types 1 and 2 are more likely to be directly adjacent to micronuclei than type 3, as would be expected when the macronuclei are derived from division of the micronuclei. The persistence of the senescent macronuclei would also explain why the MAC:MIC ratio is somewhat higher than the typical 2:1 ratio for karyorelicts.

The diversity of nuclear morphologies was one of the reasons that Foissner (1995) suggested *Kentrophoros* to be non-monophyletic. These range from a single cluster of 2 MAC + 1 MIC (e.g. *K. magnus*), to an irregular row of MAC and MIC (e.g. *K. flavus* (Xu et al. 2011)), to a nuclear capsule or complex comprising distinct MAC and MIC bounded by additional membrane structures (e.g. *K. latus* (Raikov 1972)) (Fig. 10). While the diversity of nuclei in *Kentrophoros*, in terms of number, arrangement, and now size, is striking, other karyorelict genera also show variability in their nuclear characters (Raikov 1985; Yan et al. 2017). For example, many species of *Remanella* have a single 2 MAC + 1 MIC nuclear group, with the MIC positioned between the MAC, just as in *K. magnus*, but some species, including the type *R. multinucleatus*, have an irregular row of nuclei with > 10 MAC (Xu, Gao, et al. 2013).

**Fig. 10.**
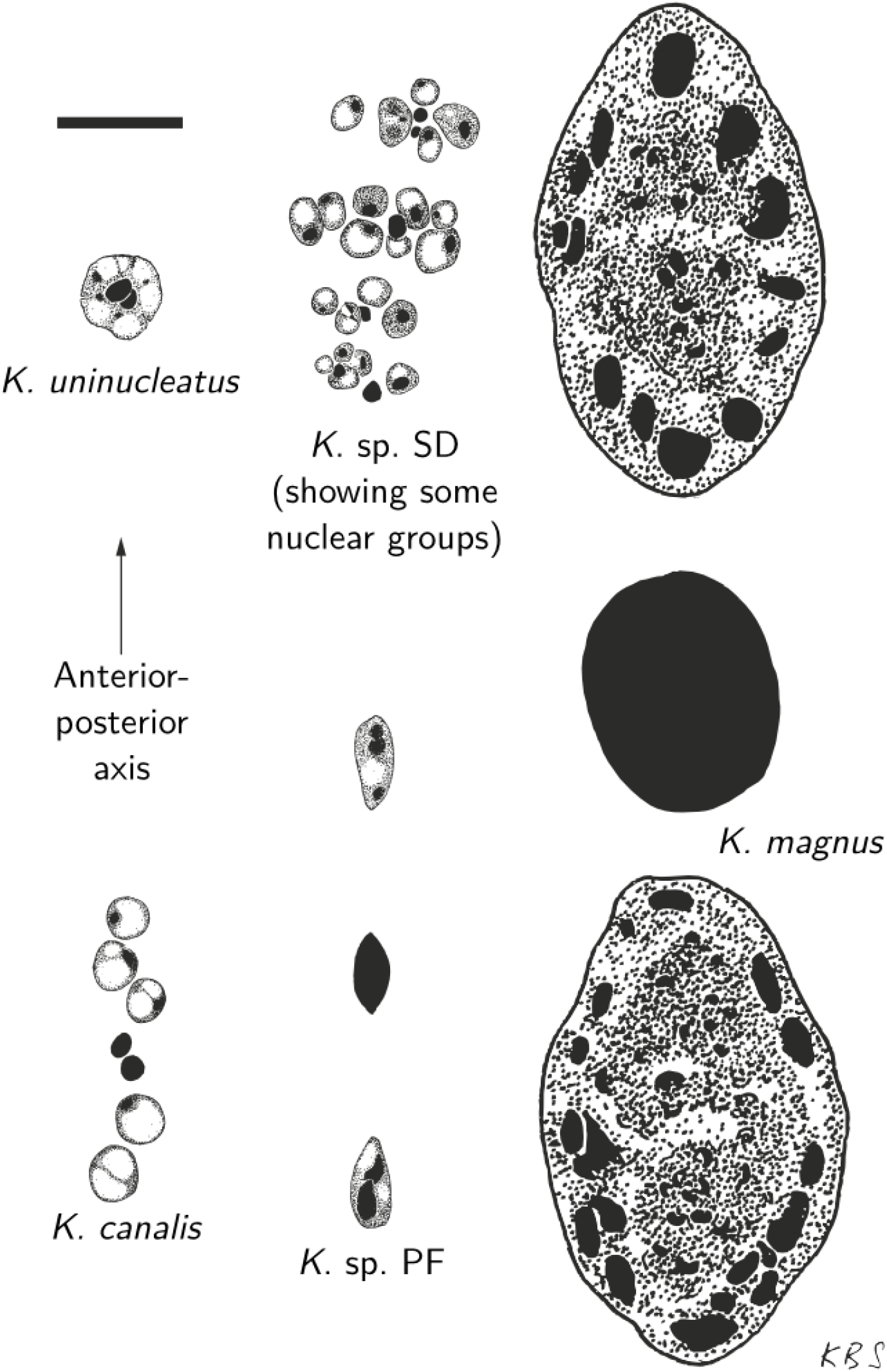
Diagrammatic drawing of nuclear diversity in *Kentrophoros* morphospecies drawn to the same scale. Scale: 10 μm.

To have a convenient shorthand for describing the nuclear arrangements of karyorelicts, we propose a “nuclear formula” (analogous to the “dental formula” used in mammalogy). If *N* is the number of nuclear groups, *x* the number of MAC, and *y* the number of MIC, then the basic formula is *N* × (*x* + *y*). If the nuclear groups are encapsulated by an additional membrane, then the formula is *N* × Cap (*x* + *y*). If the nuclei are not grouped into clusters but are in a single row aligned with the body axis, then R (*x* + *y*) where *x* and *y* are now the total number of MAC and MIC. Ambiguities and ranges of values can be indicated by question marks and hyphens respectively. For a single group of a few nuclei, distinctions between a “row” and “cluster” are probably artificial. We show how this can be applied to examples from *Kentrophoros* and other karyorelicts in Table 4.

**Table 4.** Diversity of nuclear arrangements in selected *Kentrophoros* and other karyorelicts, illustrating the use of the “nuclear formula” (see Discussion)

| Species | Formula | MAC per cluster | MIC per cluster | Type | Reference |
| --- | --- | --- | --- | --- | --- |
| *K. magnus* | R (2 + 1) | 2 | 1 | Row | This study |
| *K. canalis* | R (5 + 2) | 5 | 2 (always paired) | Row | Wright 1982 |
| *K. sp. “FM”* | R (8-40 + 4-10) | 8-40 | 4-10 | Row | This study |
| *K. gracilis* | R (7-25 + 4-21) | 7-25 | 4-21 | Row | Xu et al. 2011 |
| *K. flavus* | R (9-49 + 3-17) | 9-49 | 3-17 | Row | Xu et al. 2011 |
| *K. uninucleatus* | Cap (? + 2) | ? | 2 | Capsule | Raikov 1962 |
| *K. ponticus* | Cap (6-12 + 6) | 6-12 | 6 | Capsule | Kovaleva 1966 |
| *K. latus* | 1-4 × Cap (4 + 2) | 4 | 2 | 1-4 capsules | Raikov 1972 |
| *K. fistulosus* | ~25 × (4 + 2) | 4 | 2 | 25 clusters | Foissner 1995 |
| *Geleia sinica* | R (2 + 1) | 2 | 1 | Row | Xu et al. 2011 |
| *Remanella multinucleata* | R (7-24 + 3-9) | 7-24 | 3-9 | Row | Foissner 1996 |
| *Tracheloraphis coluber* | 1 × (4 + 2) | 4 | 2 | 1 cluster | Raikov 1985 |
| *Loxodes striatus* | 2 × (1 + 1) | 1 | 1 | 2 clusters | Raikov 1985 |
| *Tracheloraphis kahlii* | 3-11 × (6+4) | 6 | 4 | 3-11 clusters | Raikov 1985 |

### Infraciliature

Silver impregnation of karyorelicts is challenging (Foissner 2014), and unfortunately the infraciliature was poorly stained in *K.* sp. “FM”, so this discussion focuses on observations in *K. magnus*.

Foissner (1995) hypothesized that the irregularly branching fibers in the head and tail of *K. fistulosus* are homologous to fibers associated with the dorsolateral kinety of other loxodids (e.g. *Remanella*), and thus represent vestiges of the oral infraciliature. *K. magnus* also has such fibers, and like *K. fistulosus* they are found in both the anterior and posterior. If these are indeed oral vestiges, it is remarkable that they have been maintained even as other oral infraciliature structures have been lost. Given that the irregular fibers are found only in the head and tail, which are also the only symbiont-free parts of the body, they could now be performing some other function for the ciliate unrelated to food ingestion, such as sensing or movement.

The dense marginal kineties of *K. magnus* may be a previously overlooked character of some *Kentrophoros*. They are clearly visible in the head, and can be followed into the involuted mid-body region, where the dense marginal kineties on the two facing margins flank the medial suture, but they do not continue into the tail. In *K. fistulosus*, published figures show the left kinety to be flanking the medial suture of the involuted body region (Foissner (1995), Fig. 31, 45). The left kinety of *K. magnus*, however, is distinct from the dense marginal kinety and not continuous with it. In *K. flavus*, the marginal kinety of the right side is also condensed; Xu et al. (2011) state that it is condensed in the posterior region, but their figures (5J, L) show that it is condensed (compared to the other right somatic kineties) in the midbody and anterior regions too. Given that only four species (including *K. magnus*) of *Kentrophoros* have been studied with detailed attention to the infraciliature, this dense marginal kinety is probably found in other species. The irregular branching fibers (putative nematodesmata) do not appear to originate from the dense marginal kinety, so are unlikely to be homologous with the dorsolateral kinety of the loxodids.

The pair of longitudinal fibers running beside the right somatic kineties both appear to be myonemes, because they are both stained with similar intensity, and because transverse connections between fibers may connect right to left members within and between pairs. The postciliodesmata are probably not well-stained by the Fernandez-Galiano method, unlike the protargol method. Having two myonemes associated with each kinety is unlike *K. fistulosus*, which has only one myoneme on the left side of each kinety (and the postciliodesma on the right), but is more characteristic of *Geleia*. The longitudinal myonemes probably function in body contraction and movement, like in other karyorelicts and heterotrichs that have them.

The transverse connections may keep the body contractions more uniform (Lynn 2008: 126) but could also prevent herniation.

## Conclusion

*K. magnus* is a large and distinctive species, easily distinguished from known congeners by its pseudotrophosomal body form. It can therefore be considered an example of a “flagship species”, which are species that by virtue of their unusual appearance are “unlikely to be overlooked” (Foissner et al. 2008). We found *Kentrophoros* to be diverse and locally abundant in both our sampling localities (Seah et al. 2017). The fact that such species have only now been discovered points to the severe undersampling of meiofaunal ciliate diversity. Indeed, the known geographical distribution of *Kentrophoros* appears to correspond to the favored sampling spots of the handful of protistologists who have worked with karyorelicts (Table S1).

*K. magnus* may have eluded the notice of previous workers precisely because of their large size – they are easily mistaken for metazoan worms – and because of their relative rarity compared to other ciliate species. The superficial similarity to platyhelminth flatworms of the genus *Paracatenula* is particularly striking: they both have a club-shaped hyaline head/rostrum, tapered posterior tail, a pale line running down the middle of the body, and a milky appearance because of symbiotic sulfur bacteria. In our experience, the distribution of *K. magnus* is patchy; it is typical to find only tens of individuals in a bucket containing 15 liters of sediment, and often none at all. Typical extraction methods used for interstitial ciliates, such as the Uhlig method or anesthetization with magnesium chloride, use only a few milliliters of sediment at a time, and would be unlikely to recover them in significant numbers. However, meiofauna zoologists working on Elba have occasionally found these large worm-like ciliates in their collections (Giere and Erséus 2002, J. Ott, pers. comm.). We suggest that ciliatologists working with meiofauna use a variety of extraction methods (e.g. as described by Giere, 2009) as they may yield quite different results.

The shallow marine interstitial is a vast habitat, if we consider the extent of the global continental shelves. New families (Xu, Li, et al. 2013) and genera (Campello-Nunes et al. 2015) of karyorelict ciliates are still being discovered. Sampling of hitherto neglected regions, such as the Southern Hemisphere, will undoubtedly yield many more discoveries in ciliate meiofauna.

## Taxonomic summary

Class Karyorelictea Corliss, 1974

Order Protostomatida Small & Lynn, 1985

Family Kentrophoridae Jankowski, 1980

Genus *Kentrophoros* Sauerbrey, 1928

*Kentrophoros magnus* spec. nov.

### Diagnosis

Cells in vivo 2,100 ± 700 × 170 ± 23 μm (means ± standard deviations); body vermiform, involuted except for anterior (“head”) and posterior (“tail”) ends. Head and tail regions flat, hyaline, not covered by symbionts. Head club-shaped, 210 ± 45 × 77 ± 13 μm (silvered), slightly asymmetrical, with right margin more curved than left margin. Tail tapered, length 93 ± 31 μm (silvered). Anatomical margins in involuted body region meet at median suture. Left side (bearing symbionts) extensively folded and enveloped by body involution, such that bacteria apparently contained in serially-repeated pouch-like chambers (“pseudotrophosome”). Right side with ca. 100-200 longitudinal somatic diciliated kinety rows, spaced 3.0 ± 0.55 μm. Anteriormost portion of right kineties densely arranged and obliquely oriented. Marginal kineties on right side (directly flanking median suture) with densely-arranged kinetids; dense kineties extend into head and terminate before obliquely-oriented anterior kinety rows, do not extend into tail. Left side with apparently single monociliated circle kinety, but could not be traced into the mid-body region because of cell thickness and folding. Nuclei in single group at midpoint of cell, comprising two macronuclei intercalated by one micronucleus. Micronuclei oblate spheroidal, width 20 ± 2.5 μm, densely-staining, internally homogeneous. Macronuclei ovoid to teardrop-shaped, width 31 ± 4.0 μm, internally heterogeneous with numerous spheroidal chromocenters.

### Type locality

Sediment adjacent to *Posidonia oceanica* seagrass meadows off Cavoli, Isola d’Elba, Italy (42.734192 °N, 10.185868 °E, 12.8 m depth).

### Type habitat

Shallow-water marine silicate sediment, interstitial habitat.

### Type material

One holotype (NHMUK 2016.6.1.1a to o) deposited at Natural History Museum, London, and one paratype (OLML 2016/105 to 111) deposited at Oberösterreichisches Landesmuseum Linz, each comprising serial 1 μm sections of single ciliates embedded in epoxy resin, stained with toluidine blue.

### Etymology

Latin *magnus* (adj.) meaning “great” or “large”, in reference to the size of this new species compared to its congeners.

### Gene sequences

Partial 18S rRNA sequences (accession no. LT621837 to LT621844, European Nucleotide Archive) under the name “*Kentrophoros* sp. H”.

In addition, we have deposited the following voucher material for *Kentrophoros* sp. “FM”: NHMUK 2016.6.1.4 (Fernandez-Galiano silver impregnation) and NHMUK 2016.6.1.5 (SEM stub) deposited at Natural History Museum, London. OLML 2016/112 (Fernandez-Galiano silver) and OLML 2016/113 (SEM stub) deposited at Oberösterreichisches Landesmuseum, Linz. Partial 18S rRNA sequences (accession no. LT621847 to LT621850, European Nucleotide Archive) under the name “*Kentrophoros* sp. FM”.

## ACKNOWLEDGMENTS

We thank the Hydra Institute team on Elba, especially Miriam Weber, Hannah Kuhfuß, and Matthias Schneider, and the staff at Carrie Bow Caye Field Station, for invaluable support in field work. We also thank Jörg Ott for sharing personal field observations and advice, Daniela Tienken and Sten Littmann for assistance with electron microscopy, Bernd Stickfort for library support, Brenda Seah for Russian translations, Monika Bright for helpful suggestions and arranging support for JMV, and the Core Facility of Cell Imaging and Ultrastructure of at the University of Vienna. Financial support was provided by the Max Planck Society to BS, HG, and ND, and the Gordon and Betty Moore Foundation grant GBMF 3811 to ND. JMV was supported by Austrian Science Fund (FWF) grant P24565-B22 to Monika Bright.

## Author contributions

BKBS designed study with HRGV and ND, collected specimens, performed experiments, analyzed data, and wrote paper with HRGV and ND. JMV performed TEM of *K. magnus*. NL performed SEM of *K.* sp. “FM”. TS prepared holotype of *K. magnus*.

---

**Table S1.** Known collection localities of *Kentrophoros*

| **Locality** | **Reference** |
| --- | --- |
| Fetovaia, Elba, Italy | Seah et al., 2017 |
| Cavoli, Elba, Italy | Seah et al., 2017 |
| Sant' Andrea, Elba, Italy | Seah et al., 2017 |
| Twin Cayes, Belize | Seah et al., 2017 |
| Sagami Bay, Japan | Takishita et al. 2010 |
| Sagami Bay, Japan | Takishita et al. 2010 |
| Helgoland, Germany | Kahl 1935 |
| Sylt, Germany | Seah, unpubl. data |
| Kiel, Germany | Sauerbrey 1928 |
| No. 1 Beach, Qingdao, China | Gao et al. 2010; Xu et al. 2011 |
| Roscoff, France | Fauré-Fremiet 1950; Foissner 1995 |
| Barnstable, MA, USA | Fauré-Fremiet 1951 |
| Swansea Bay, UK | Wright 1982 |
| Niva Bay, Denmark | Fenchel and Finlay 1989 |
| Gullmaren, Sweden | Hedin 1977 |
| Lemon Bay, FL, USA | Noland 1937 |
| Dal'niye Zelentsy, Barents Sea | Raikov 1974 |
| Wattenmeer, Germany | Wickham, Gieseke, and Berninger 2000 |
| Gryaznaya Bay, White Sea | Burkovsky and Mazei 2010 |
| Baie Omega, Sevastopol | Raikov 1971 |
| Jubail, Saudi Arabia | AL-Rasheid 1998 |
| Banyuls-sur-Mer, France | Dragesco 1954a |
| Ussuri Bay, Russia | Raikov 1963 |
| Posyet, Russia | Raikov and Kovaleva 1968 |
| Astafyev Bay, Russia | Raikov and Kovaleva 1996 |

